## Supplementary material for "Integrating and formatting biomedical data as pre-calculated knowledge graph embeddings in the Bioteque": Supp Information

### Supplementary Figures and Tables

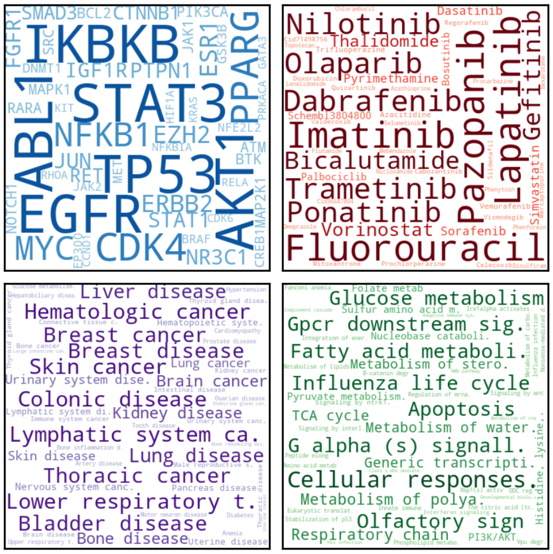


**Supplementary Figure 1. Node popularity in the propagated graph.** Most popular nodes in the KG within the gene (GEN, blue), compound (CPD, red), disease (DIS, purple) and pathway (PWY, green) universe. In contrast to Fig. 1F, dataset associations were propagated across the corresponding ontologies (when possible) before computing the popularity of the nodes.

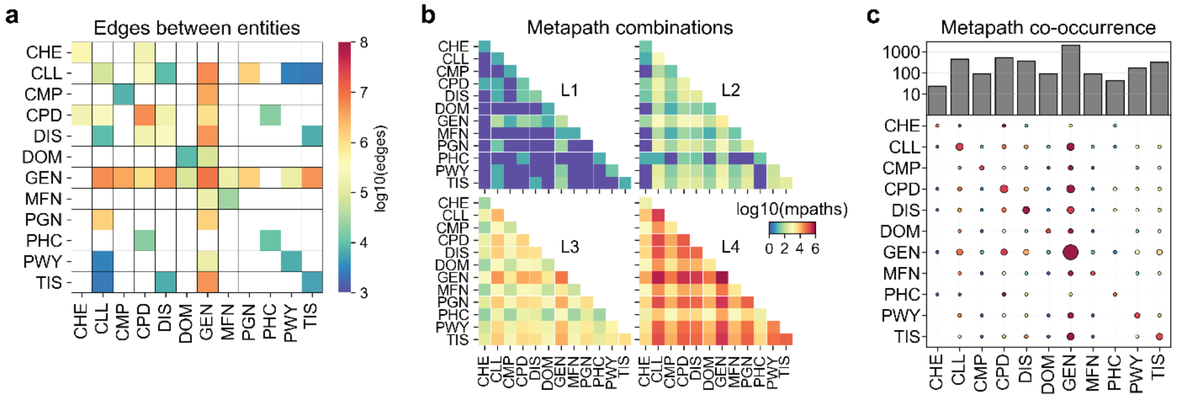


**Supplementary Figure 2. Number of relations in the KG. a** Number of possible edges between entities considering all associations and datasets available in the graph. **b** Theoretical number of metapaths of length 1, 2, 3, and 4. **c** Number of times that every entity (x-axis) co-occurs in a metapath with another entity (y-axis). The colourr scale illustrates the most used entities in each row, red and blue being the most and less used entities, respectively. The size is proportional to the number of times a given entity participates in a metapath.

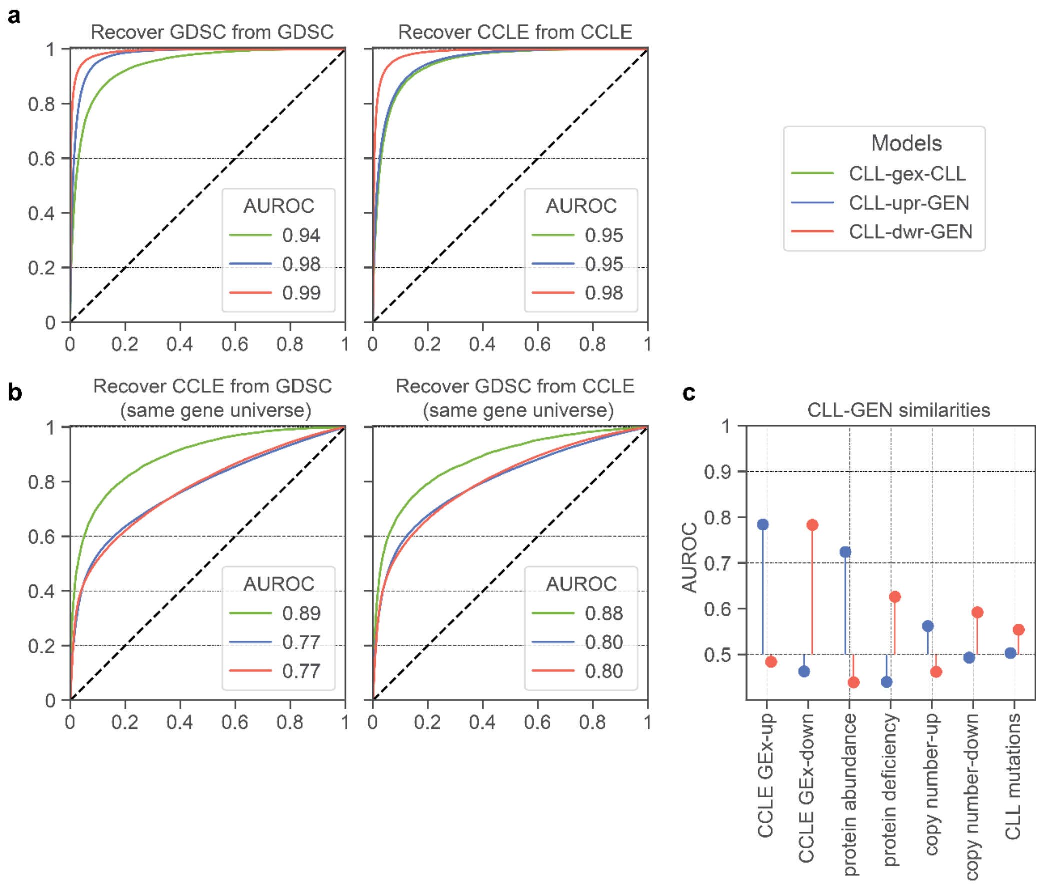


**Supplementary Figure 3. Agreement between GDSC and CCLE embeddings. a** Recovery of the original GDSC (left) and CCLE (right) network by their corresponding Bioteque embeddings. **b** Recovery of the CCLE (left) and GDSC (right) panels using the GDSC embeddings and CCLE embeddings, respectively. In contrast to Fig. 5D, embeddings were obtained from a GDSC and CCLE version in which we kept only those genes in common between both panels. **c** Characterization of the cell-gene (CLL-GEN) similarities for the 'cell upregulates gene' (CLL-upr-GEN*)* and 'cell downregulates gene' (CLL-dwr-GEN*)* metapath embeddings.

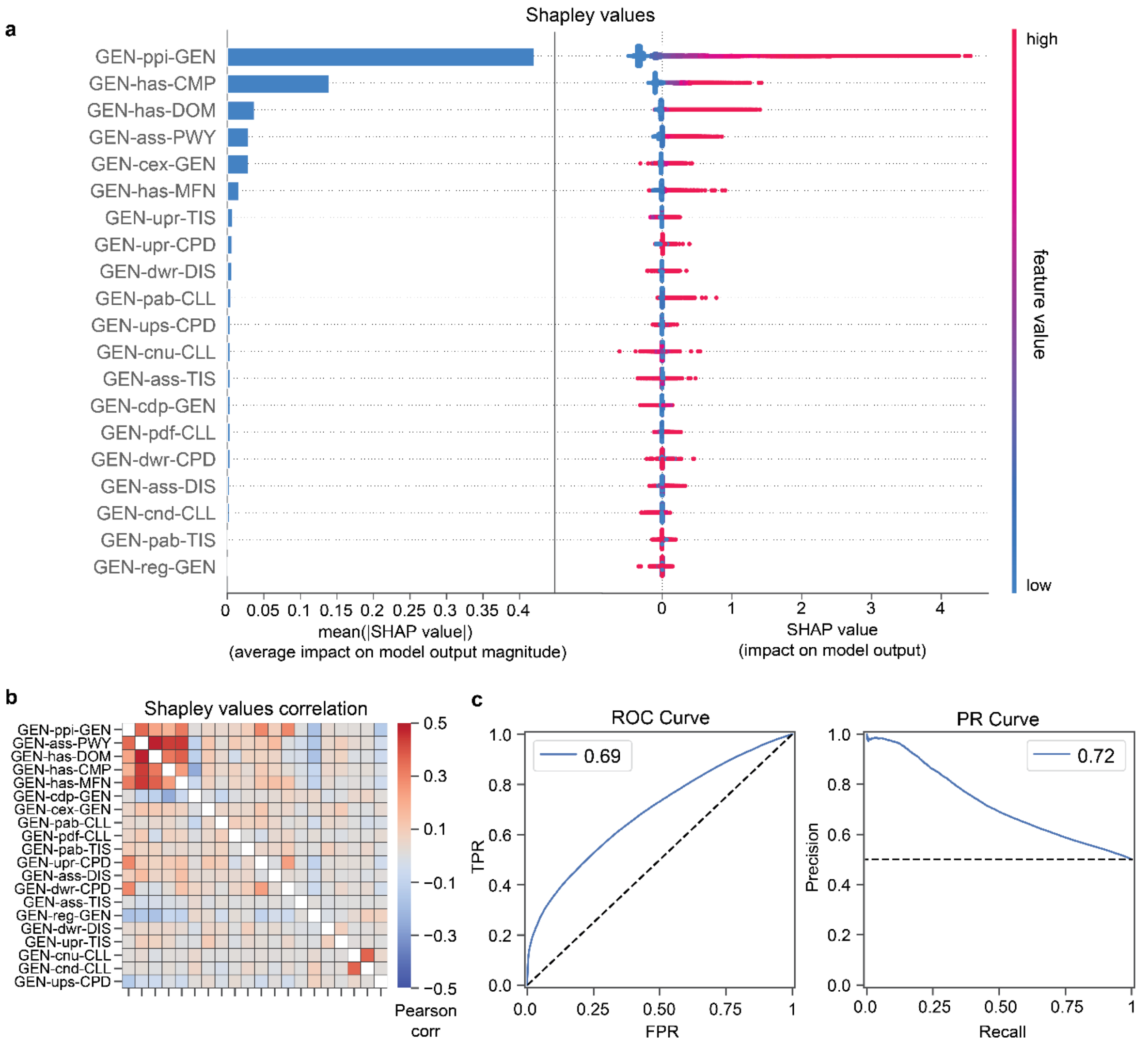


**Supplementary Figure 4. Metapath importance when predicting HuRI-III PPIs. a** Feature importance measured as Shapley values (x-axis) for each metapath (y-axis) when predicting HuRI-III PPIs. The higher the Shapley value the higher the impact when predicting the correct class. (Left) Mean absolute Shapley value for each metapath. (Right) Individual Shapley values for each prediction (i.e., each dot corresponds to a predicted PPI). In the colour scale, red and blue indicate lower and higher P-values for the corresponding metapath, respectively. **b** Pairwise Pearson correlation matrix of the Shapley value vectors for each metapath. The higher the correlation, the higher the agreement between metapaths when classifying a PPI. **c** ROC (left) and Precision-Recall (right) curves obtained from the model. A precision of 0.5 is expected at recall 1 as negative pairs were downsampled to achieve a 1:1 balance with positive pairs.

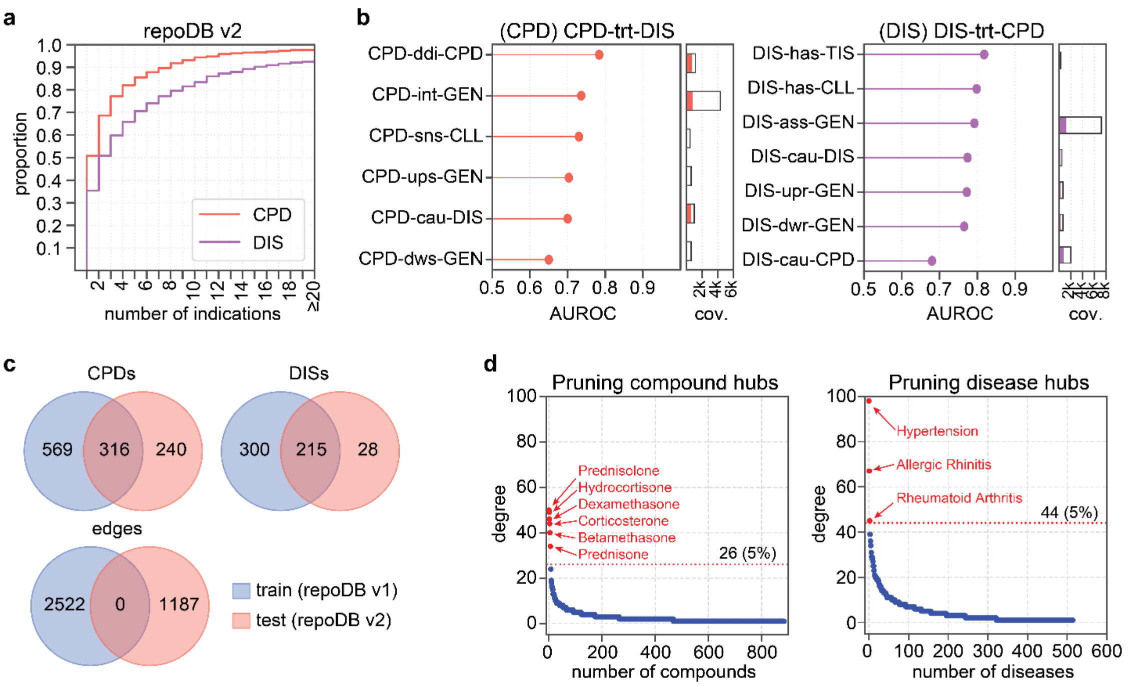


**Supplementary Figure 5. Exploring the repoDB dataset. a** Cumulative distribution of the number of treatment indications reported by repoDB (v2) for the compounds (CPD, red) and diseases (DIS, purple). **b** Top L1 metapaths recapitulating CPD-CPD (left) and DIS-DIS (right) treatment similarities. This was measured by assessing (AUROC) how the different embeddings up-ranked CPD-CPD (or DIS-DIS) pairs associated with the same treatment. The bar plots show the number of nodes available by each metapath, colouring those that were covered by the 'compound treats disease' (CPD-trt-DIS) network**. c** Number of unique compounds (CPDs), diseases (DISs) and edges in the train (repoDB v1) and test (repoDB v2) splits after mapping the entities to the 'compound interacts protein' (CPD-int-GEN) and 'disease associates with gene' (DIS-ass-GEN) embedding universes. **d** Compounds (left) and diseases (right) of the train split were sorted according to their node degree in the repoDB CPD-trt-DIS network. The red line shows the degree corresponding to 5% of total possible associations. We highlight those compounds and diseases whose degree exceeded this limit and were, therefore, pruned.

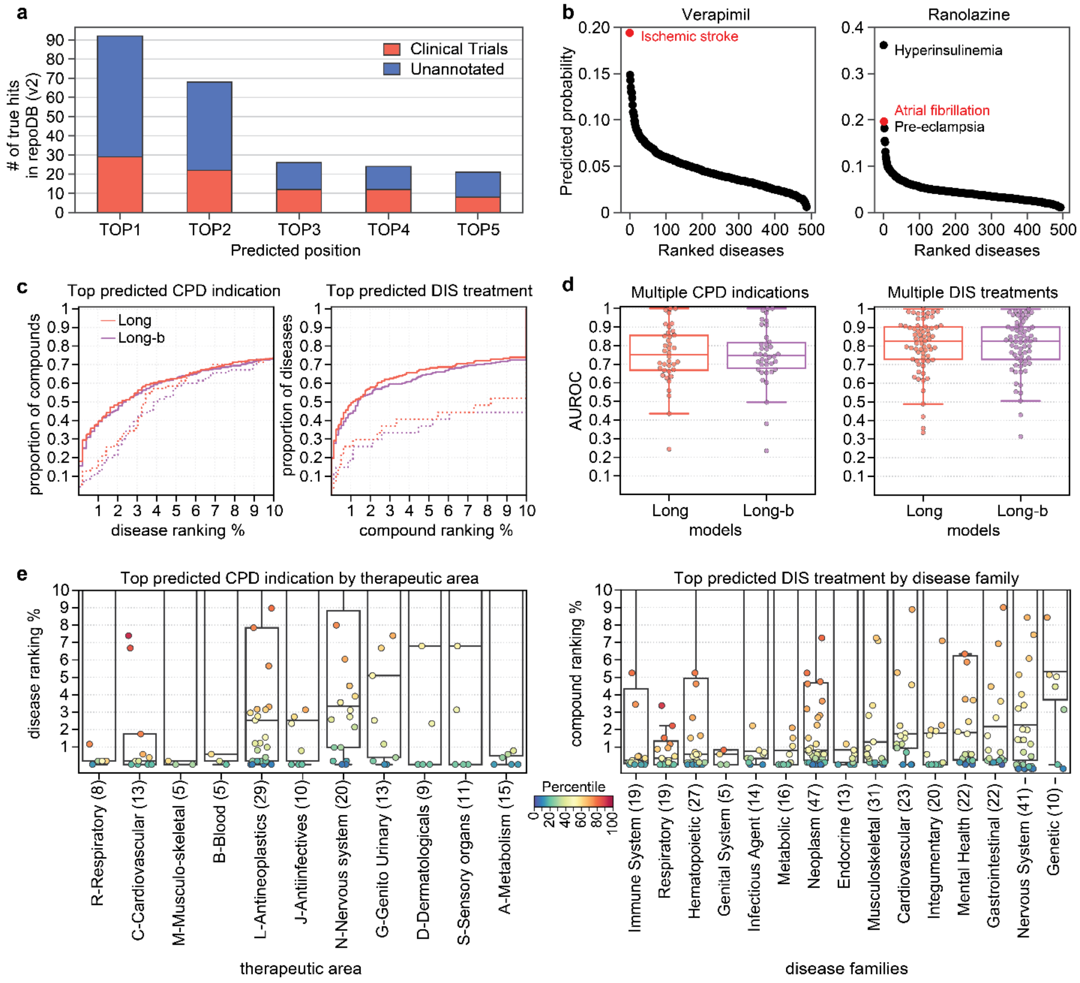


**Supplementary Figure 6. Additional repoDB prediction results. a** Number of compound-disease (CPD-DIS) repurposing pairs from repoDB (v2) (y-axis) correctly predicted by the *Long* model within the top 5 positions (x-axis). In red we colour those predictions for which repoDB provides evidence of having been in clinical trials. **b** Predictions for all the screened new indications for Verapamil (left) and Ranolazine (right) drugs ranked (x-axis) according to the predicted probability given by the *Long* model (y-axis). In red we show those new indications validated in repoDB (v2). **c** Cumulative distribution (y-axis) of compounds (left) and diseases (right) according to the ranked position (x-axis) of the best predicted disease indication (left) or compound treatment (right) for the *Long* and *Long-b* models. The rankings are shown in percentages and only for the first 10% of compounds or disease predictions. Dotted line shows the distribution for those compounds or diseases with only one positive indication in repoDB (v1). **d** Classification performance obtained for each compound (n=38, left plot) and disease (n=67, right plot) with multiple (>=5) new indications reported in repoDB (v2). Box plots indicate median (middle line), 25th, 75th percentile (box), and max and min value within the 1.5*25th and 1.5*75th percentile range (whiskers). **e** We categorized the compounds (left) and diseases (right) according to their therapeutic area and disease family (x-axis) and showed the ranking of the best predicted indication and treatment (y-axis), respectively. Each dot corresponds to either a drug or a disease. The parenthesis in the x-axis indicates the total number of drugs or diseases in each class. Box plots indicate median (middle line), 25th, 75th percentile (box), and max and min value within the 1.5*25th and 1.5*75th percentile range (whiskers). Since the ranking was cut at the closest 10% of predictions, we coloured each drug or disease by the percentile it represents in the population of the corresponding group.

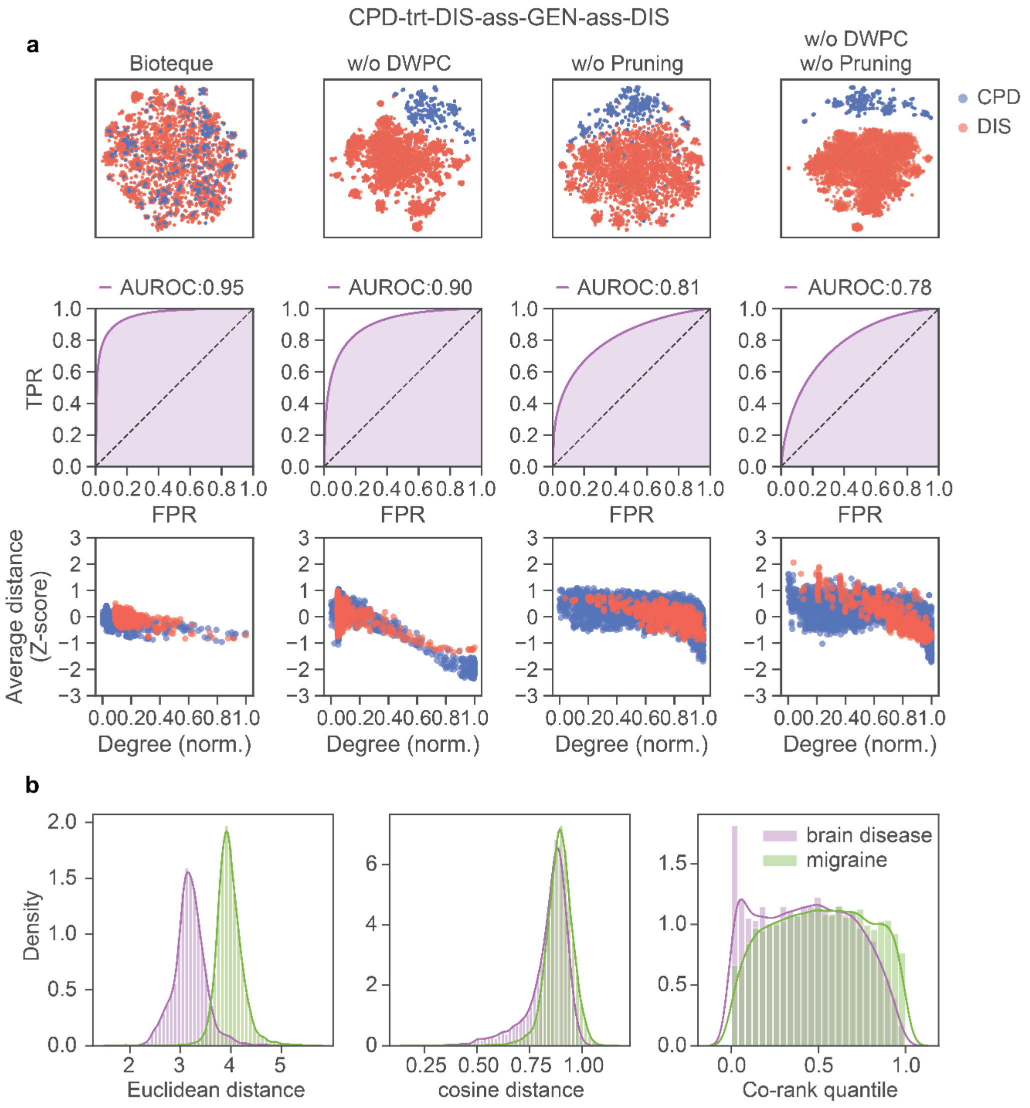


**Supplementary Figure 7. Accounting for node degree biases. a** Using the metapath embedding CPD-trt-DIS-ass-GEN-ass-DIS as reference we calculated 3 other embedding spaces where we removed the DWPC weights (w/o DWPC), the limitation in the number of edges (w/o pruning) or both (w/o DWPC, w/o pruning). Looking at the 2D representations (first row) we can see how removing either the DWPC or the pruning introduces biases in the space according to the node type, making them cluster separately in the 2D projection and affecting the ability of the space to recapitulate the original KG (second row). In the last row, we show the association between the average z-score cosine distance of each node (y-axis) and their normalized degree (i.e., divided by the max degree within each node type) in the network (x-axis). Notice that, while it is expected that nodes with a higher degree will be, by definition, closer to more nodes, the average distance does not differ more than 1 standard deviation from the average (z-scores between -1 and 1). However, removing either the DWPC or the pruning makes higher-degree nodes much closer, on average, to any other node in the space. **b** Distance distribution of the ‘Brain disease’ and ‘Migraine’ nodes to each of the genes available in the GEN-ass-DIS embedding space (obtained from DisGeNET). From left to right we show the distribution using Euclidean distances, cosine distances (1-cosine similarity), and Co-rank quantiles (calculated as specified in the Methods section).

**Supplementary Table 1. Comparing pre-existing knowledge graphs.** **Although our KG includes up to 150 datasets, we selected 66 as a reference to perform the embeddings*

| KG dataset | Design Use case | Entities | Edges | Entity Types | Relation Types | Contains Features | Datasets | Version info | Last update |
|---|---|---|---|---|---|---|---|---|---|
| Hetionet | Repurp. | 47K | 2.2M | 11 | 24 | no | 29 | no | 2017 |
| DRKG | Repurp. | 97K | 5.7M | 13 | 107 | yes | 34 | no | 2020 |
| BioKG | General | 105K | 2M | 10 | 17 | yes | 13 | no | 2020 |
| PharmKG | Repurp. | 7.6K | 500K | 3 | 29 | yes | 7 | no | 2020 |
| OpenBioLink | Benchmark | 184K | 4.7M | 7 | 30 | no | 17 | no | 2020 |
| Clinical KG | Personalized medicine | 16M | 220M | 35 | 57 | no | 35 | no | 2020 |
| Bioteque (ours) | General | 450k | 30M | 12 | 67 | no | 150 (66*) | yes | 2022 |
